# p53-dependent polyploidisation after DNA damage in G2 phase

**DOI:** 10.1101/2020.06.09.141770

**Authors:** Anna Middleton, Rakesh Suman, Peter O’Toole, Karen Akopyan, Arne Lindqvist

## Abstract

Cell cycle progression in the presence of damaged DNA can lead to accumulation of mutations and pose a risk for tumour development. In response to DNA damage in G2 phase, human cells can be forced to exit the cell cycle in a p53-p21- and APC/C^Cdh1^-dependent manner. Cells that exit the cell cycle in G2 phase become senescent, but it is unclear what determines this commitment and whether other cell fates occur. We find that a subset of immortalised RPE-1 cells and primary human fibroblasts spontaneously initiate DNA re-replication several days after forced cell cycle exit in G2 phase. By combining single cell tracking for more than a week with quantitative immunofluorescence, we find that the resulting polyploid cells contain increased levels of damaged DNA and frequently exit the cell cycle again in the next G2 phase. Subsequently, these cells either enter senescence or commit to another round of DNA re-replication, further increasing the ploidy. At least a subset of the polyploid cells show abnormal centrosome numbers or localisation. In conclusion, cells that are forced to exit the cell cycle in G2 phase face multiple choices that lead to various phenotypes, including propagation of cells with different ploidies. Our findings suggest a mechanism by which p53-positive cells can evade senescence that risks genome integrity.

**Main points:**

-Cell cycle exit from G2 phase does not necessarily lead to senescence
-Resumption of proliferation after G2 phase cell cycle exit starts with DNA replication
-Successive cell cycle exits lead to propagation of cells with different ploidies
-A p53-dependent mechanism allows eventual proliferation after DNA damage

## Introduction

Cell cycle progression in the presence of damaged DNA can lead to propagation of mutations. In response to damaged DNA, cells pause the cell cycle, in particular prior to initiating DNA replication or cell division. This pause, termed a checkpoint, is poorly sustained over time. In case of excessive or sustained DNA damage, cells therefore permanently block proliferation, either by regulated cell death, or by entering into senescence. In G2 phase, the road to senescence is initiated by p53- and p21-dependent activation of APC/C^Cdh1^^1–5^. Activation of APC/C^Cdh1^ leads to degradation of many key regulators of G2 phase and mitosis, thereby re-setting the cell cycle. Whereas we have referred to this phenomenon as cell cycle exit^6^, others described it as cell cycle withdrawal^7^ or mitosis skip^8^. Common to all is the description that this is the first irreversible step leading to senescence from G2 phase. While it can be modulated by checkpoint duration^8^, efficiency of DNA repair^9^, and CDK activity in G2 phase^10^, the irreversible step rendering cell cycle exit independent of upstream checkpoint kinase signalling is marked by translocation of Cyclin B1 to the nucleus^6–8^.

While cell cycle exit occurs rapidly in single cells, establishment of senescence is a gradual process involving large changes in gene expression, chromatin rearrangements into stress foci, and establishment of a secretory programme^11–13^. How cell cycle exit is maintained until senescence is firmly established remains unclear. Yet, the regulation of this step is likely important during tumour development. Activation of oncogenes commonly leads to replication stress, resulting in S phase cells containing damaged DNA. The resulting DNA damage response (DDR) eventually leads to induction of senescence and the formation of a preneoplastic lesion^14^. To develop into a neoplastic lesion, at least some cells need to evade senescence^15,16^.

Cell growth continues during a DNA damage response, and the changed ratio between DNA and cell volume contributes to senescence^17^. One way to restore this ratio is to increase the amount of DNA by increasing cell ploidy. A number of mechanisms can contribute to the formation of large polyploid cells. Except cell fusion, any type of cessation of the cell cycle between DNA replication and physical cell division, followed by a cell cycle restart will increase ploidy^18^. In the absence of p53, cells containing DNA damage can eventually enter mitosis, which risks mitotic exit before cytokinesis ^1,19^. Alternatively, during sustained DNA damage in G2 phase, p53-negative cells can resume DNA replication without entering mitosis^20^. Combined with the ability of p53 to block polyploid cells in G1 phase, p53 is therefore considered as a guardian against the formation of polyploidy^21–23^.

We have studied the long-term cell fate decisions after cell cycle exit from G2 phase in RPE-1 cells and primary human fibroblasts. Using fluorescent live-cell imaging in combination with quantitative phase imaging, we report a mechanism of polyploidisation that paradoxically depends on p53. We find that a subpopulation of cells can re-enter the cell cycle and initiate re-replication ~ 4 days after DNA damage-induced cell cycle exit in G2 phase. The resumed replication is associated with DNA damage, which in the following G2 phase results in cells either entering mitosis or exiting the cell cycle again. Repeated rounds of cell cycle exit followed by senescence or resumed proliferation create a heterogenous population with various ploidies, cell sizes, and proliferative potentials. Taken together, p53-dependent cell cycle exit in G2 phase can be reversible and does not solely lead cells to senescence.

## Results

### A setup to study DNA damage-induced cell cycle exit in G2 phase

To create a system in which cells robustly exit the cell cycle in G2 phase, we synchronised hTERT RPE-1 cells in G2 phase and induced DNA damage by addition of Etoposide. For efficient synchronisation, we added Hydroxyurea (HU) to block cells in S-phase. After release from HU-synchronisation, cells progressed through G2 phase to mitosis, indicating that lasting DNA damage due to HU-mediated nucleotide depletion was limited (Figure S1). Since the majority of cells were in late S phase 7 hours after synchronisation release, we selected this time point to induce DNA damage in further studies (Figure 1a). After addition of Etoposide to the synchronous cells, we used fluorescent live-cell imaging to assess cell cycle exit in G2 phase. Initially, cells did not enter mitosis after Etoposide addition, indicating that a checkpoint was firmly established (Figure 1b). After several hours of checkpoint arrest, cells showed translocation of Cyclin B1-YFP to the nucleus, followed by a loss of Cyclin B1-YFP with no sign of mitosis, characteristic of cell cycle exit from G2 phase^6–8^ (Figure 1c). Similarly, Cyclin A2-eYFP was degraded as the control cells entered mitosis and after Etoposide addition Cyclin A2-eYFP levels were initially sustained, indicating a checkpoint-arrest in G2 phase. After a delay, Cyclin A2-eYFP expression declined, indicating that cell cycle exit had occurred (Figure 1d). Given the essential role that p21-dependent APC/C^cdh1^ activation and subsequent degradation of cell cycle proteins play in cell cycle exit in G2 phase, we last monitored APC/C^cdh1^ activity by expressing RFP conjugated to the APC/C degron motif from geminin. Geminin-RFP levels started to decrease after 6 hours of Etoposide treatment, but stabilised in the presence of the APC/C inhibitor ProTAME, indicating that APC/C was activated (Figure 1e). In line with the live-cell data, immunofluorescent staining suggested that the majority of G2 phase cells had exited the cell cycle, since there was a loss of Cyclin A2 and increased p21 expression (Figure 1f). Taken together these data indicate that cell cycle exit in G2 phase can be robustly established within 24 hours of Etoposide treatment.

**Figure 1.**
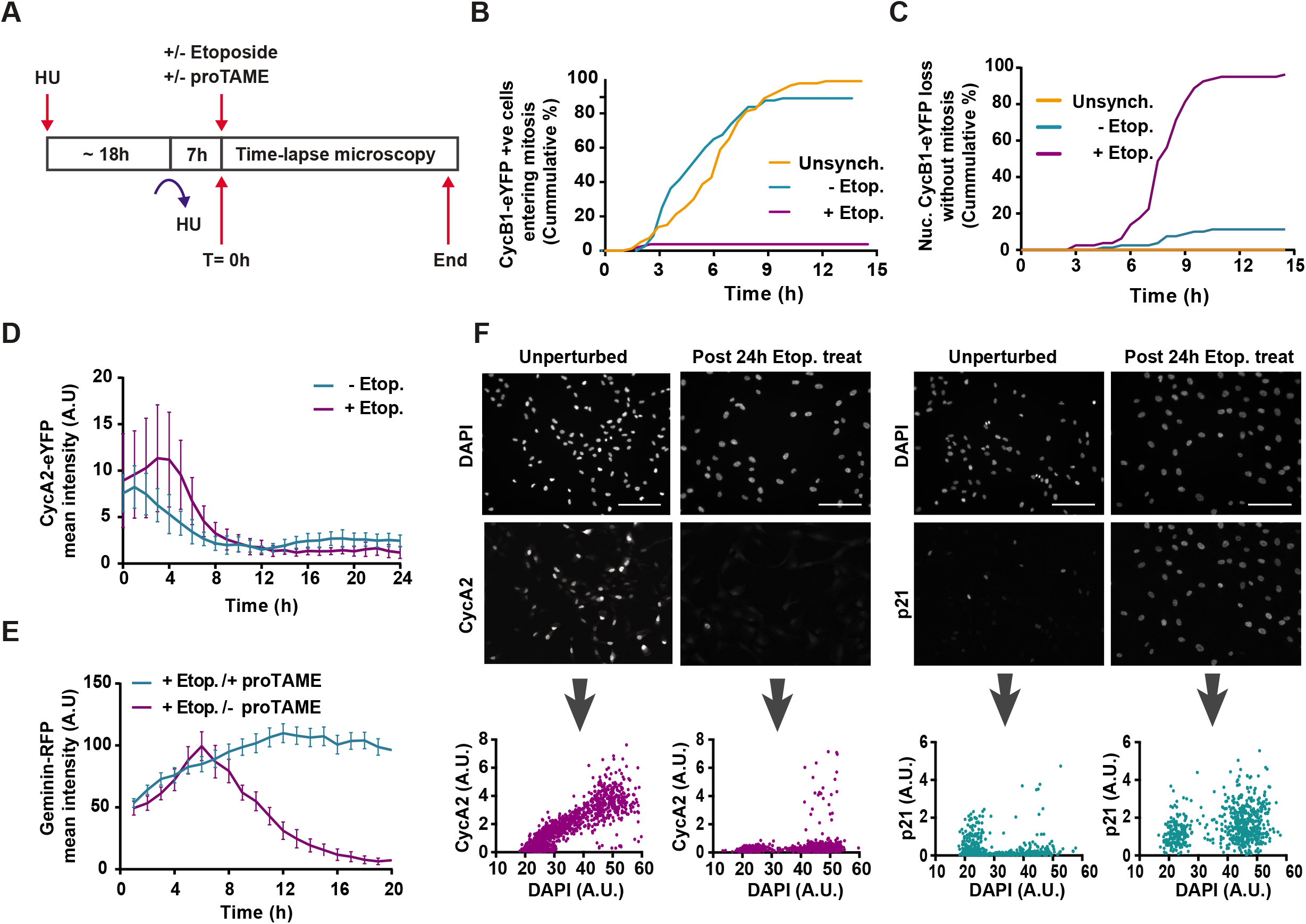
A setup for Etoposide-induced cell cycle exit in G2 phase. (a) Schematic showing the experimental setup. 7h after hydroxyurea synchronisation release, cells were treated either with or without 2μM Etoposide or 15μM ProTAME. (b and c) RPE-1 Cyclin B1-eYFP cells. Data from images acquired at 30min intervals. (b) Cumulative % of Cyclin B1-eYFP positive cells entering mitosis (n= 80 cells per condition). (c) Cumulative % of Cyclin B1-eYFP positive cells showing nuclear Cyclin B1-eYFP translocation and degradation without entering mitosis (n=80 cells per condition). (d) RPE-1 Cyclin A2-eYFP cells. Quantification of general Cyclin A2-eYFP expression. Data from images acquired at 1h intervals (n=10 positions; Mean image intensity ± SD). (e) RPE-1 cells expressing Geminin-RFP. Quantification of general Geminin-RFP expression. Data from images acquired at 1h intervals (n= 10 positions; Mean image intensity ± SEM). (f) Representative images and quantification of immunofluorescent staining for Cyclin A2 (n=656 cells per condition) and p21 (n=1551 cells per condition), and DAPI staining for nucleus. Scale bar: 2mm.

### A subset of cells become polyploid after G2 phase cell cycle exit

Previously we and others have reported that cell cycle exit from G2 phase can lead to senescence^7,8,10^. However, whether senescence is a strict consequence of cell cycle exit in G2 phase is unclear. Since a key characteristic of senescence is the inability of cells to resume the cell cycle, we decided to assess long-term cell fate after G2 phase cell cycle exit. To identify cells that would exit the cell cycle from G2 phase, we pulse-labelled with EdU before inducing DNA damage (Figure 2a and 2b). As expected, the majority of EdU-positive cells contained 4n DNA-contents 24 hours after addition of Etoposide (Figure 2c, upper left). Six days after Etoposide wash out, the majority of EdU positive cells still contained 4n DNA content, indicating that a majority of cells did not resume proliferation. Importantly, we did not detect EdU-positive 2n cells, suggesting that cell cycle resumption from G2 phase did not occur (Figure 2c, lower left). However, we found that a subset of EdU-positive cells became polyploid 6 days after Etoposide wash out, suggesting that G2 exited cells could resume at least some aspects of the cell cycle.

**Figure 2.**
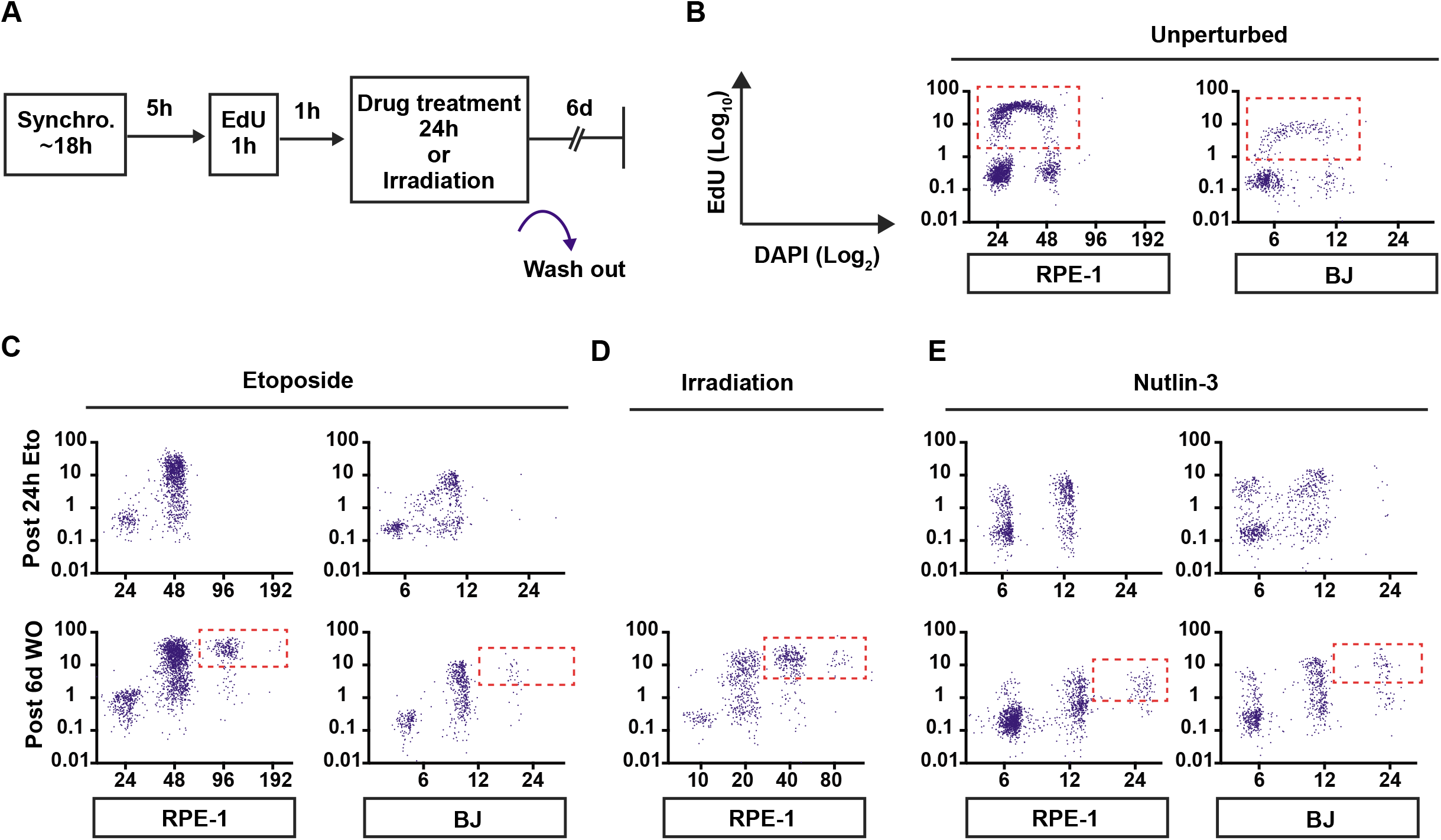
Cell cycle exit in G2 phase leads to polyploidisation. (a) HU synchronised RPE-1 and BJ cells were pulse labelled with EdU before 24h drug treatment (2μM Etoposide; 12μM Nutlin-3) or γ-irradiation and fixed for DAPI and EdU staining after 6 days. (b) Unperturbed cells (RPE-1 n= 2017; BJ n=629). (c) After 24h Etoposide treatment (Upper panel: RPE-1 n=1154 cells; BJ n=625 cells). 6 days after Etoposide wash out (Lower panel: RPE-1 n=2012 cells; BJ n=644 cells). (d) 7 days after 16 gray γ-irradiation in RPE-1 cells (n=845 cells). (e) 24h after Nutlin-3 treatment (Upper panel: RPE-1 n=850 cells; BJ n=760 cells) and 6 days after Nutlin-3 wash out (Lower panel: RPE-1 n=1532 cells; BJ n=855 cells). EdU positive polyploid cells are marked by a red-dotted box.

To test if the observed polyploidy depended on treatment with Etoposide, we monitored DNA content after γ-irradiation. Similar to after Etoposide treatment, populations of irradiated EdU-positive cells contained polyploid DNA contents, but not 2n DNA contents, showing that polyploidisation is not restricted to Etoposide (Figure 2d). To separate whether polyploidisation depends on DNA damage itself or on DNA-damage induced cell cycle exit, we treated cells with Nutlin-3, which can induce cell cycle exit in G2 phase in the absence of exogenous DNA damage by interfering with Mdm2-directed p53 degradation^7^. EdU-positive cells accumulated in a 4n population after 24hours of Nutlin-3 treatment, indicating that cell cycle exit in G2 phase had occurred, albeit less efficiently than by DNA damage insult, as a notable number of EdU-positive 2n cells were also present (Figure 2e, upper left). Importantly, the presence of EdU-positive polyploid cells 6 days after Nutlin-3 wash out suggests that polyploidisation is not due to DNA damage per se, but is rather a consequence of p53-dependent cell cycle exit in G2 phase (Figure 2e, lower left). Given that RPE-1 cells are immortalised cells, we sought to test if polyploidy can occur in a primary cell line. Although primary human foreskin fibroblasts were less responsive to HU for synchronisation, the DNA-contents profile of EdU positive cells after 6 days of either Etoposide or Nutlin-3 wash out were comparable to RPE-1 cells (Figure 2c and 2e, right). This suggests that polyploidisation is not limited to immortalised RPE-1 cells.

### Long-term live-cell imaging reveals heterogeneity in cell fate after cell cycle exit in G2 phase

To gain further insight into how cells become polyploid after cell cycle exit in G2 phase, we sought to monitor cell cycle signalling in individual cells following Etoposide-induced DNA damage. Since Cdk activity is the key driver of cell cycle progression, and is instrumental to initiate DNA replication, we monitored RPE-1 cells expressing a Cdk activity sensor. The Cdk activity sensor localises in the nucleus when Cyclin E-Cdk2 and Cyclin A-Cdk1/2 are inactive and gradually translocates to the cytoplasm as activity increases when the cell cycle progresses^24,25^. We followed cells by combined live-cell fluorescence microscopy and quantitative phase imaging (QPI), allowing simultaneous detection of Cdk activity and cell dry mass^26,27^. As polyploidy was not observed until several days after DNA damage (Figure 2), we sought to track single cells for more than a week. To allow for cell tracking while minimising phototoxicity, we acquired QPI images every 10 minutes and fluorescence images every 30 minutes. Unperturbed cells went through successive rounds of cell division, as expected showing cytoplasmic Cdk sensor before mitosis and an increase in nuclear Cdk sensor after mitosis (Figure 3e and Movie S1). After each division, the integrated QPI intensity that is proportional to the total dry mass of the cell decreased, whereas the average QPI intensity that depends on cell morphology increased sharply as cells rounded up during cell division^26^. At late stages in the experiment, individual cells were difficult to segment and quantify accurately due to crowding, further highlighting that the experimental setup allowed long-term proliferation (Figure 3e and Movie S1). Before crowding, both average and integrated QPI intensities fluctuated around similar levels, suggesting that cell mass and morphology did not change over time due to the imaging conditions. Thus, the imaging setup can be used to follow cells for more than a week.

**Figure 3.**
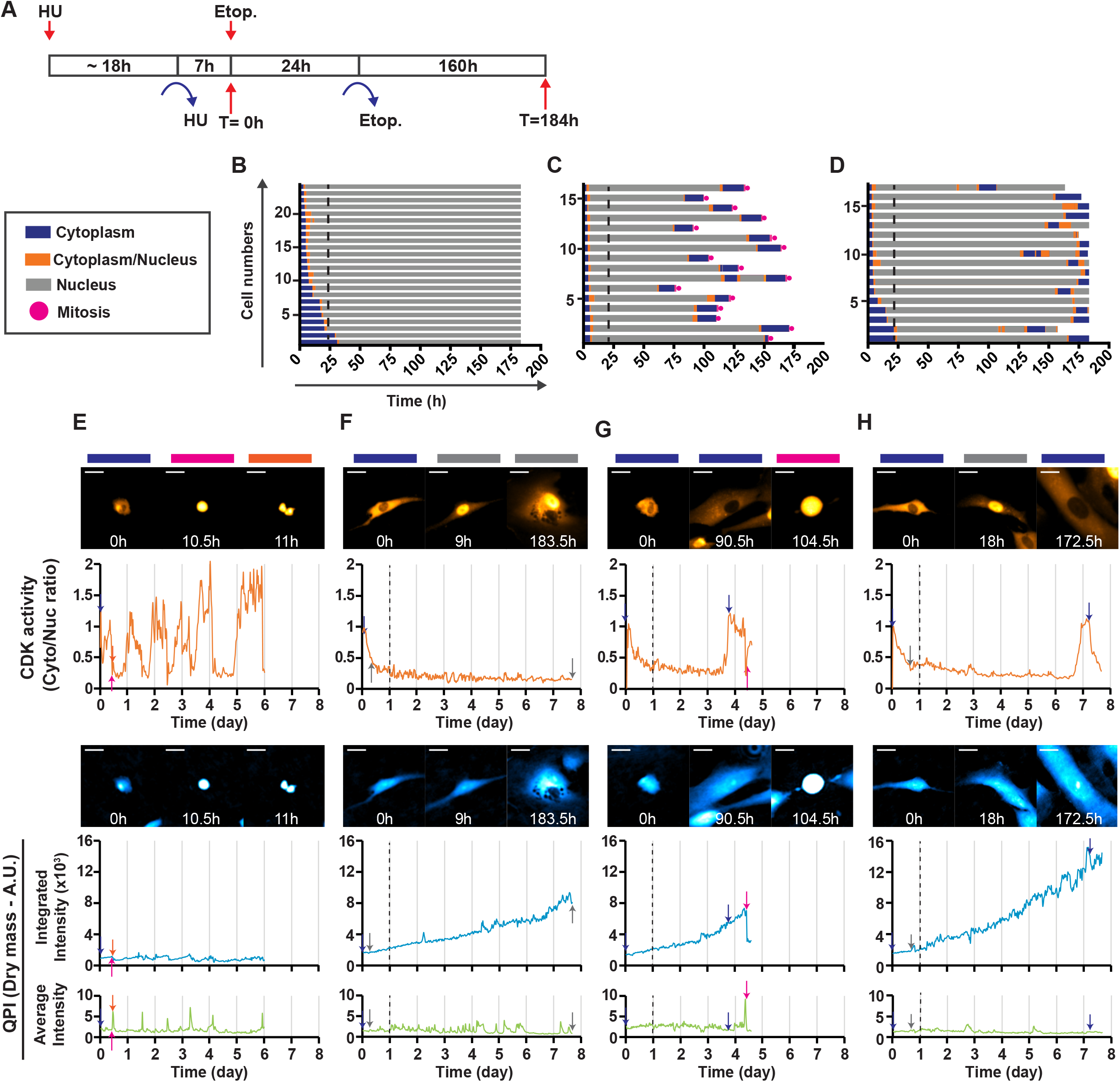
Long-term time-lapse imaging of RPE-1 cells with Cdk-RFP sensors reveals multiple cell fate decisions after exit in G2 phase. (a) HU synchronised RPE-1 cells expressing Cdk-RFP sensors were followed by fluorescence and quantitative phase (QPI) time-lapse imaging for more than 7 days at 30 min and 10 min intervals, respectively. (b-d) Each horizontal bar represents the Cdk localisation trace of a single cell over time. The black dotted line indicates the time of Etoposide wash out. There are three main groups based on localisation of the Cdk sensor after initial nuclear translocation and cell cycle progression in response to Etoposide treatment (b) no localisation changes, (c) translocation to the cytoplasm followed by mitosis and (d) translocation to cytoplasm with no progression to mitosis. (e-h) Quantification of Cdk-sensor localisation (top), integrated QPI intensity (middle), and average QPI intensity (bottom) of a single cell. (e) Unperturbed cell cycle. (f-h) Cells treated with Etoposide. A representation of each of the three groups based on Cdk activity and cell cycle progression after initial Cdk activity loss (f) no Cdk re-activation, (g) Cdk re-activation followed by mitosis and (h) Cdk re-activation with no progression to mitosis. Time of the represented images are indicated on the image’s respective graphs with colour arrows. Scale bar: 30μm. Please note that to provide a continuous read-out, the Cdk sensor reading is at an arbitrary low value in mitosis.

After 24h treatment with Etoposide, proliferation was largely blocked. Strikingly, cells occupied a larger surface area and the integrated QPI intensity continuously rose, indicating that the cell mass of individual cells increased up to 10 times during the course of the experiment (Figure 3f-h). In contrast, the average QPI intensity of cells fluctuated over time, reflecting morphology changes during cell migration. However, the overall levels were largely similar to control cells, indicating that whereas the cells became larger, they retained a similar dry mass density (Figure 3f-h and Movie S1). The Cdk sensor translocated to the nucleus in most cells within 24 hours of Etoposide treatment (Figure 3b-d and Movie S1). Etoposide did not impair Cdk activity in S phase, as the Cdk sensor showing partial cytoplasmic expression did not immediately translocate to the nucleus. However, as cells entered G2 phase, the fluorescence intensity ratio showed that Cdk activity reduced rapidly and persisted for a few hours at a low level before activity was lost completely (Figure 3f-h). This Cdk dynamic is comparable to our previous study^10^, indicating that cell cycle exit had occurred in G2 phase after DDR activation.

To exclusively monitor cells treated in G2 phase, we selected cells expressing the Cdk sensor in the cytoplasm at the time of Etoposide treatment. We observed that Cdk dynamics varies from cell to cell after 6 days of Etoposide removal. In order to understand cell fate decisions, we categorised cells into three groups based on their Cdk activity and cell cycle progression after wash-out of Etoposide. The first group that we identified were cells that never regained Cdk activity, but showed a gradual growth in dry cell mass, suggesting that cell cycle exit had led to a permanently arrested state (Figure 3b and 3f). Apart from this group, we observed that Cdk sensors translocated back to the cytoplasm in approximately 50% of the traced cells. Approximately 50% of these cells underwent cell division, giving rise to daughter cells that contained half the dry mass, but were still larger than unperturbed cells (Figure 3c and 3g). The last group represents the other half of cells that regained Cdk activity but did not progress to mitosis. Similar to the response after Etoposide addition, these cells lost Cdk activity again without entering mitosis (Figure 3d and 3h). Interestingly, in both groups, the duration and pattern of regained Cdk activity was similar to the duration and pattern of Cdk activity during S and G2 phases during unperturbed proliferation. This shows that the pulse of Cdk activity could be of sufficient duration to encompass DNA replication. Further, our data shows that large polyploid cells with single large oval nuclei observed in Figure 2 do not stem from cytokinesis failure, endomitosis, or cell fusion (Movie S1). Given that cell cycle exit in G2 phase depends on APC/C^Cdh1^ activation, which during unperturbed growth mediates the progression from mitosis to G1 by targeting G2/M regulators for degradation, cells that exited the cell cycle in G2 phase are likely in a similar biochemical state to G1 phase. We propose that resumption of Cdk activity after DNA damage induced cell cycle exit in G2 phase leads to DNA re-replication and polyploidy. Furthermore, after the initial Cdk re-activation, occasional cells again both lost and re-gained Cdk activity (Figure 3c and 3d). Such repeated activation could in principle account for additional increases of ploidy beyond 8n.

Since we observed three major groups of cell fates after Etoposide treatment, we assessed if the timing of cell cycle exit after induction of DNA damage is a determining factor for cell fate. We sorted cells in order of when the Cdk sensor was inactivated after Etoposide addition and compared that with cell fate. There were no correlations between Cdk inactivation timing and cell fate in group 1 and 3 (Figure 3b and 3d). However, cells that progressed to mitosis lost Cdk activity within 12 hours of Etoposide addition (Figure 3c). The timing of cell cycle exit depends on cell cycle position when Etoposide is added^6^. This suggests that predominately cells that were in late G2 phase at the time of DNA damage entered mitosis after cell cycle resumption. Taken together, these results demonstrate that p53 positive cells several days after Etoposide treatment in G2 phase can re-initiate S phase upon cell cycle resumption and are able to complete the cell cycle.

### G2 phase exited cells re-enter the cell cycle with unresolved DNA damage in the presence of DDR

To gain insight into DNA damage signalling in cell populations that re-initiate Cdk activity, we used a fixed-cell quantitative immunofluorescence approach (Figure 4a). As the Cdk sensor responds to Cyclin A- or Cyclin E-dependent Cdk activity^20,24,25^, we selected Cyclin A2 to identify the cell population that regains Cdk activity (Figure 4b). Quantification of DAPI and Cyclin A2 staining in cells that were pulse labelled with EdU, revealed that Cyclin A2 levels were low in 4n cells, indicating sustained cell cycle exit from G2 phase. However, Cyclin A2 levels progressively built up as DNA contents increased to 8n. This Cyclin A2 expression pattern resembled that of an unperturbed cell cycle with a shifted ploidy, suggesting that resumed Cyclin A expression is coupled to re-initiated DNA replication. However, the majority of 8n cells did not express Cyclin A2, which was different from unperturbed 4n cells that contain high Cyclin A2 levels (Figure 4b). One reason for this difference could be that Cyclin A2 negative cells exit the cell cycle again, similar to the initial exit after Etoposide treatment. This would be consistent with live cell data, in which Cdk activity persisted for a period sufficient to complete S phase before this Cdk activity was again lost (Figure 3d). We therefore sought to assess whether polyploid cells contain damaged DNA by determining γH2AX levels as a marker for active DDR kinases (Figure 4c). Total γH2AX intensity levels were low in unperturbed cells and showed the highest level during DNA replication. As expected, the γH2AX level was increased in response to Etoposide treatment. The increase in γH2AX level was slightly reduced 6 days after Etoposide wash out, possibly reflecting ongoing DNA repair. 8n cells contained higher γH2AX levels than 4n cells. However, this difference in γH2AX staining largely disappeared when correcting for DNA content (Figure 4c and f). Thus, with respect to Cyclin A2 levels, cells regaining Cdk activity resemble unperturbed cells, but with a shifted ploidy and increased γH2AX levels.

**Figure 4.**
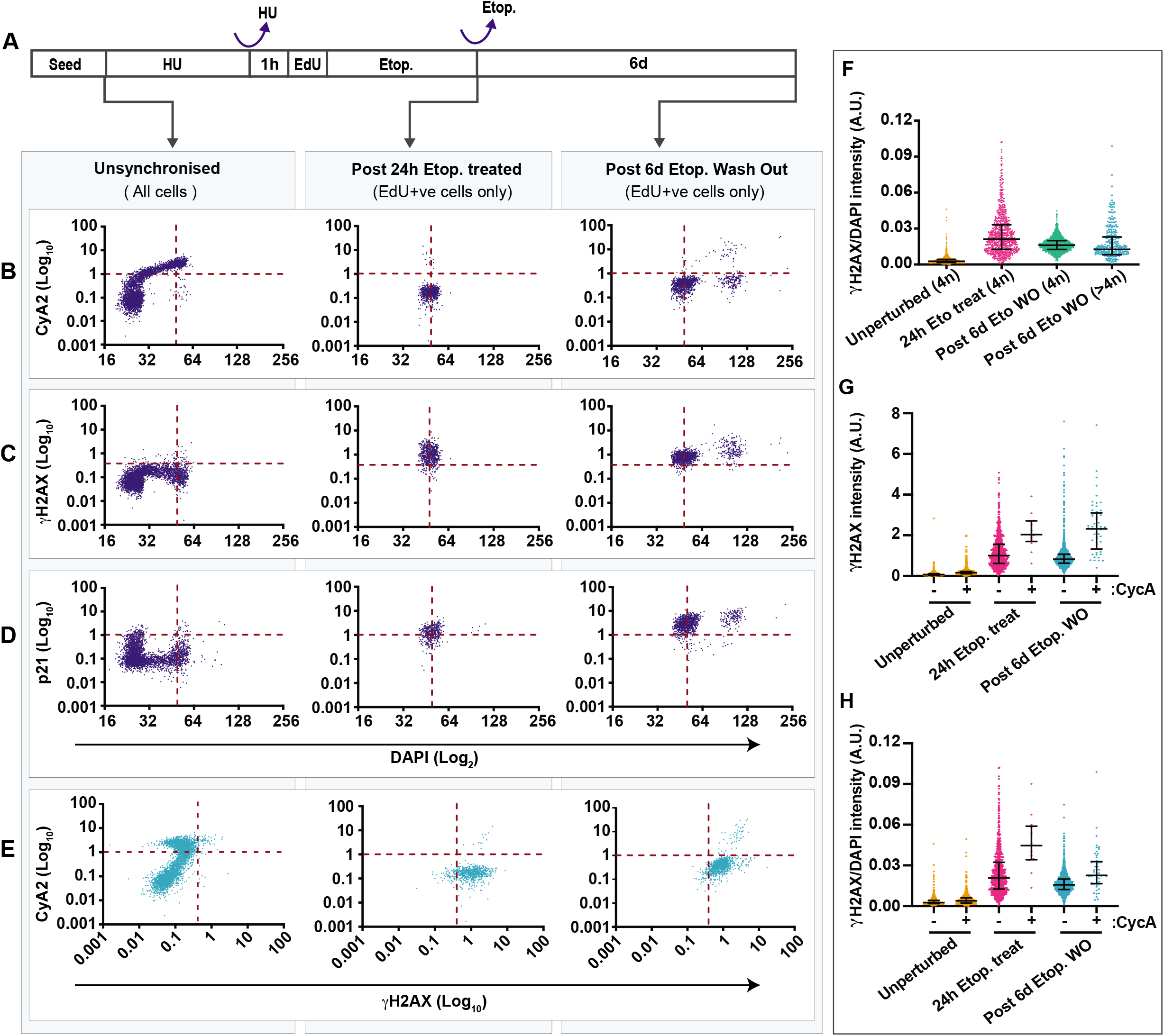
G2 phase exited cells resumed the cell cycle with unresolved DNA damage and in the presence of DDR. (a) Schematic showing the experiment setup. Quantification of immunofluorescence of (b) nuclear Cyclin A, (c) mean nuclear γH2AX and (d) nuclear p21 intensity versus DNA contents by DAPI staining intensity in single RPE-1 cells. Only EdU-positive cells are represented in the graphs except under unsynchronised conditions. The vertical red dotted line indicates G2 phase cells, while the horizontal red dotted line marks the base line for positivity of the indicated proteins. (e) Quantification of immunofluorescence of nuclear Cyclin A2 levels versus nuclear mean γH2AX intensity in single RPE-1 cells. (f) Mean γH2AX intensity of single-cells divided by their DAPI signal.

To assess if γH2AX increased after re-initiation of Cdk activity, we co-stained cells with antibodies against γH2AX and Cyclin A2 (Figure 4e). Interestingly, Cyclin A2 positive populations exhibited higher γH2AX levels, indicating that either cells with high γH2AX reinitiate proliferation, or more likely, that Cdk activity and associated DNA replication cause additional DNA damage. If DNA damage contributes to cell cycle exit, this should lead to significant p21 induction. p21 expression was low in the unperturbed cells, particularly during S phase, which reflects PCNA/DNA replication mediated degradation of p21 (Figure 4d). After 24 hours of Etoposide treatment, the time when most cells had exited the cell cycle, p21 levels were markedly increased. During 6 days after Etoposide removal, p21 levels increased further, indicating sustained DDR signalling. However, similar to the γH2AX levels, there was a spread in p21 levels among individual cells. A subset of cells showed lower p21 levels, in particular cells with ploidies between 4n and 8n, suggesting engagement in DNA replication. 8n cells showed increased p21 and γH2AX, which reflects DDR activation upon completion of DNA replication. Taken together, these data indicate that cells regaining Cyclin A expression also re-initiate DNA replication, which in turn leads to a reduction in p21. However, as cells progress to an 8n G2 phase, they contain active DNA damage signalling and start to re-accumulate p21.

### Cell cycle re-entry can lead to re-replication of centrosomes

During the cell cycle, duplication of DNA is frequently coordinated with duplication of centrosomes^28^. Since we observed re-initiation of DNA replication after cell cycle exit, we sought to investigate whether centrosomes were also re-duplicated. Using immunofluorescence, we visualised the subcellular localisation of the centrosomal component pericentrin. As expected, pericentrin predominantly appeared in one or two foci near the nuclear envelope (Figure 5a). We then used the setup described in Figure 4a and identified EdU positive polyploid cells. Although the majority of polyploid cells contained an unchanged number of pericentrin foci, some EdU positive polyploid cells contained more than two foci, or pericentrin foci that were displaced from the nuclear envelope (Figure 5a). As direct counting of centrosomes was impractical in a high-content approach, as a proxy we instead quantified the levels of pericentrin staining in centrosomes (Figure 5b). In unperturbed cells, pericentrin intensity increased through S phase to be approximately doubled in G2 phase. Etoposide treated 4n cells over time showed similar intensities as unperturbed G2 phase cells, which indicates that pericentrin intensity is not affected by cell cycle exit. Interestingly, although the distributions overlap, EdU-positive polyploid cells showed increased pericentrin levels. Taken together, these data suggest that centrosome re-duplication can occur in cells that eventually regain Cdk activity after cell cycle exit in G2 phase.

**Figure 5.**
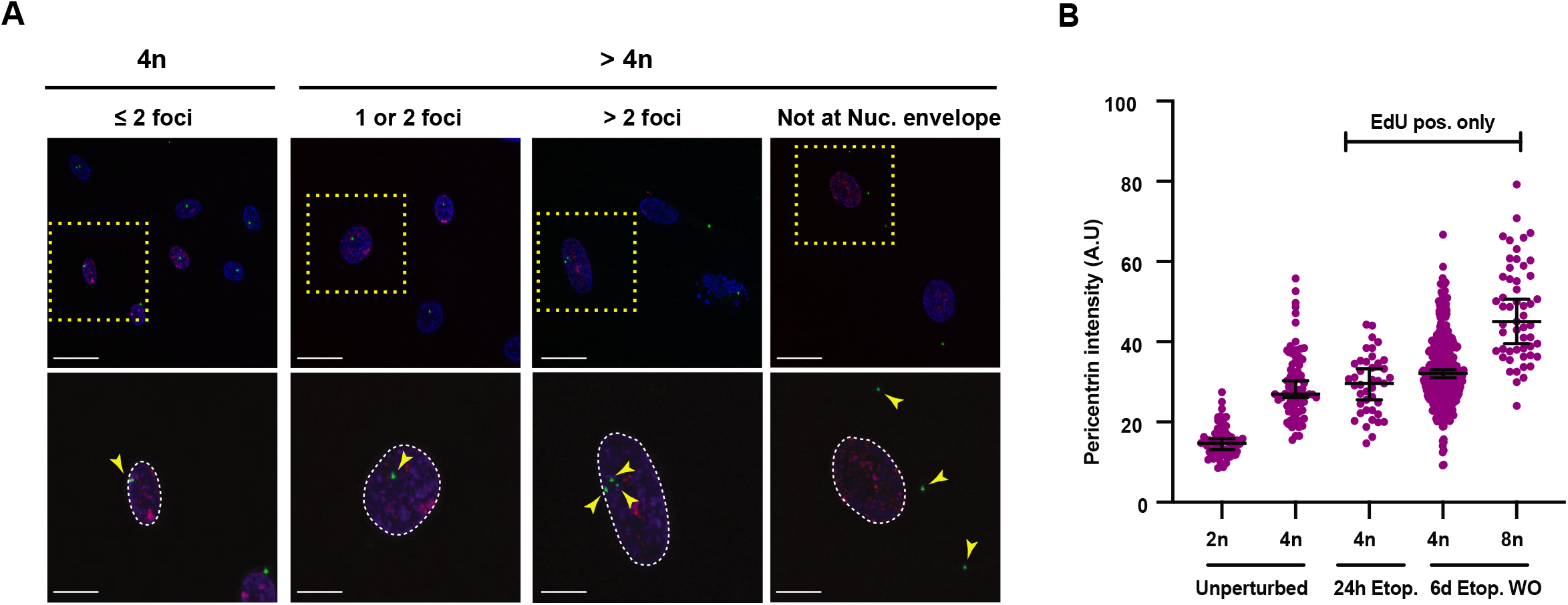
Cell cycle re-entry leads to increased pericentrin intensity and localisation abnormalities. Post 7h HU released RPE-1 cells were pulse-labelled with EdU prior to 2μM Etoposide treatment and fixed for DAPI (blue), EdU (red) and anti-pericentrin antibody (green) staining after the indicated time points. (a) Representative maximum intensity projections of deconvolved z-stacks. The yellow dotted boxes indicate the corresponding enlarged image below. The location of Pericentrin foci are indicated by yellow arrows. (b) Immunofluorescence quantification of Pericentrin antibody staining. Only EdU-positive cells are represented in the graph, except under unsynchronised conditions. Unperturbed cells (n=64; 2n cells, n=74; 4n cells). Post 24h Etoposide treatment (n=41 cells). Post 6d Etoposide wash out (n=314; 4n cells, n=53; >4n cells). Graph shows median intensity with interquartile range for each ploidy. Scale bar:50μm (Upper row), 20μm (Lower row).

## Discussion

Polyploidy and aneuploidy are common features in different types of cancer, contributing to cancer progression, surveillance escape and relapse (Reviewed in^29,30^). Polyploidy does not always arise from bona fide re-replication, and other mechanisms have been reported-cell fusion, endomitosis and acytokinetic mitosis (Reviewed in^31^). Our work has extended these findings by studying cell fate after DNA damage-induced cell cycle exit in G2 phase in untransformed cells. Previous efforts to determine cell fate decisions after cell cycle exit in G2 have monitored proliferation or senescent states at fixed time points in pooled-population or by short-term observations^7,8,32^. By monitoring Cdk activity in single-cells, we discovered that cell cycle exit in G2 phase is not a permanently arrested state and cells do not necessarily progress to senescence. Importantly, our long-term observations have allowed us to follow these infrequent cell cycle re-entry events over time, revealing that cells initiate DNA re-replication upon cell cycle resumption, rather than recovering from a biochemical G2 state. In addition, our single-cell data showed that the cell fate of polyploid cells diverges again in the following G2 phase towards cell division or a renewed cell cycle exit, suggesting a mechanism for creating a heterogenous population of proliferating and senescent cells with various ploidies.

What determines re-activation of Cdk in a subpopulation of cells after G2 phase exit remains elusive, although expression of Cdk inhibitors are likely to contribute. The memories of variable mitogen and stress signals convert into a competition between Cyclin D1 and p21 expression that contribute to regulate the proliferation-quiescence decisions in daughter cells^33–36^. Similarly, variable p16 expression could lead to different cell fate decisions, as it also stoichiometrically inhibits Cyclin D1-Cdk4/6^37^, and upregulation of Cyclin D2 contributes to proliferation of p53-positive polyploid cells^38^. In the proliferation-arrest decision circuit, oscillation of p53 triggers a sharp switch between p21 and Cdk2 allowing cells to escape from arrest^39^. In addition, Cdk activity during G2 phase stimulates p21 production to enforce cell cycle exit^10^. Thus, the dynamics of Cdk inhibitors may be established at different levels in individual cells both before and after cell cycle exit from G2 phase, suggesting a possible mechanism of why a subset of cells escape from cell cycle arrest in the presence of DNA damage and intact functional p53.

Our work reveals a novel mechanism of polyploidisation that paradoxically depends on p53. P53-dependent premature activation of APC/C^Cdh1^ leads to degradation of mitotic regulators such as Cyclin A and Cyclin B, thereby lowering Cdk activity to allow licensing of replication. Importantly, APC/C^Cdh1^ also targets geminin, whose depletion is sufficient to induce re-replication in the presence of functional p53^40^. This suggests the resetting of G2 phase cell cycle configuration to G0/G1 phase, as a consequence of p53-mediated cell cycle exit, allows initiation of re-replication.

It may come as somewhat of a surprise that polyploid cells either progress to mitosis and complete the cell cycle, or repeat a cell cycle exit in G2 phase. It is unclear to what extent the repeated cell cycle exit depends on DNA damage and a reinforced DDR gained during re-replication. Parts of the DDR play key roles in regulating the cell cycle in the absence of external DNA damage, in which DNA replication itself activates ATR/Chk1 (reviewed in ^41,42^). Furthermore, cells recovering from quiescence were recently shown to be susceptible to replication stress^43^. Stable homologous recombination repair intermediates efficiently sustain ATR/Chk1 signalling, leading to an enhanced checkpoint and eventual cell cycle exit in G2 phase^9^. In mice, disruption of Fen1-PCNA interaction leads to replication stress and polyploidy in a Chk1-dependent manner^44^. This suggests that during replication after cell cycle re-entry, replication errors, including homologous recombination repair intermediates, may trigger a second cell cycle exit in the following G2 phase. Interestingly, polyploid tumour cells in particular are known to rewire their DNA damage response and repair pathways to allow proliferation after replication stress^45^. Thus, it is reasonable to think that additional DNA damage and fitness of the DDR network might be a contributing factor for determining cell fate commitment after re-replication.

Extra centrosomes can generate chromosome instability by promoting asymmetric chromosome segregation^46^. We show that at least a subpopulation of polyploid cells can gain extra centrosomes during re-replication. This opens up the possibility that polyploid cells could give rise to aneuploidy through chromosome mis-segregation, which could increase genetic heterogeneity and thereby contribute to tumour evolution. Nonetheless, re-replication and depolyploidisation has been suggested as a strategy for cancer cells to escape chemotherapy. Several types of cancer cells can re-replicate in response to Etoposide treatment^47,48^. Also, microenvironment changes similar to those in solid tumours induce polyploidisation in transformed and non-transformed cell lines, mainly driven by re-replication^49^. Diploid cells that arise from polyploid tumour cells have been shown to gain chemotherapy resistance^47,50^. Similarly, regular-sized tumours derived from polyploid giant cancer cells also have an increased resistance to chemotherapy^51,52^. In this regard, polyploid cells with the capability to proliferate could be implicated in the development of drug resistance.

Our study revealed that cell cycle exit in G2 phase leads to various cell fates, including re-replication and proliferation, which create heterogenous cell populations with various ploidies. While cell cycle re-entry after G2 phase exit is relevant to tumourigenesis in terms of genomic instability, the question remains as to whether cells have an indefinite proliferative potential. With regards to cell cycle regulation, future studies would need to investigate the mechanisms that determine cell fate after cell cycle exit.

## Materials and Methods

### Cell culture

Human hTERT-RPE-1 cells (a kind gift from René Medema), hTERT-RPE-1 Cyclin B1-YFP^53^, and hTERT-RPE-1 Cyclin A2-eYFP^54^ were cultured with DMEM/F-12 + GlutaMax (Life Technologies), and BJ primary human foreskin fibroblasts (a kind gift from Bennie Lemmens) were cultured in DMEM, in an ambient-controlled incubator at 37°C with 5% CO_2_. All cells were supplemented with 10% foetal bovine serum (FBS; HyClone) and 1% Penicillin/Streptomycin (Pen/Strep; Gibco), unless stated otherwise.

### Cell synchronisation

Cells were synchronised by overnight incubation with 2mM Hydroxyurea (HU; sigma-Aldrich). HU was dissolved in the cell culture media and filtered at each time of use. Synchronised cells were washed three times with PBS before being released into fresh media.

### Flow cytometry assay

Cells were seeded in a 6-well imaging plate (Cornings) at 60% confluency. Cells were harvested by trypsinisation and washed with PBS before fixing with −20°C 70% ethanol (Sigma-Aldrich). PBS/0.1% tween-20 was used to wash the cells, which were resuspended in PBS for DNA staining with propidium iodide (35uM). Cells were analysed by FACSCanto II (Becton Dickinson) and the data was processed with BD FACSDiva (Version 8.0) software.

### Immunostaining

Immunofluorescent staining was performed as described previously^6^. The following antibodies were used in this study: Cyclin B1 V152 (4135, Cell Signalling), Cyclin A2 H432 (SC-751, Santa Cruz Biotechnology), γH2AX (9781, Cell Signalling), p21 Waf1/Cip1(12D1) (2947, Cell Signalling), pericentrin (ab4448, Abcam), Alexa Fluor 488-Goat anti-rabbit (4412, Cell Signalling), Alexa Fluor 647-Goat anti-rabbit (A21245, Invitrogen), Alexa Fluor 647-Donkey anti-mouse (715-605-150, Jackson ImmunoResearch), Alexa Fluor 488-Goat anti-rabbit (A11034, Life technologies) and Alexa Fluor 488-Donkey anti-mouse (715-545-150, Jackson ImmunoResearch).

### Live-cell microscopy and quantification

Cells were seeded in a 96-well imaging plate (BD Falcon) at 40% confluency. The growth media was replaced with 10% FBS and 1% Pen/Strep supplemented Leibovitz’s L-15 medium (Invitrogen), 2h before the start of imaging. Live-cell images were acquired at 37°C using an ImageXpress system (Molecular Devices) with a 20x objective (0.45 NA). Images were processed and analysed using ImageJ software. For figure 1b and c, cells were scored manually. Cell division was scored as the last time point before cytokinesis. For figure 1d and e, images were corrected by subtracting the background using a rolling ball algorithm. The average fluorescence intensity per image was quantified and the average background intensity, estimated from areas without cells, was subtracted.

### Fixed-cell microscopy and quantification

Cells were grown in a 96-well imaging plate (BD Falcon) and fixed with 3.7% formaldehyde (Sigma Aldrich) for 5 min followed by permeabilization in ice-cold methanol for 2 min. Images were acquired using an Operetta (PerkinElmer) 20x(0.45 NA) objective. Images were corrected by subtracting the background using a rolling ball algorithm on ImageJ software. To measure the nuclear fluorescence intensity of single-cells, the images were screened for single-cell selection semi-manually using CellProfiler software. Cell cycle stages were determined based on mean intensity of DAPI levels. To subtract different backgrounds for individual cells, a background function was applied to the mean nuclear fluorescence intensity. For Pericentrin analysis, a DeltaVision Spectris imaging system (Applied Precision), with 20x objective (0.75 NA) and an ImageXpress system (Molecular Devices) with a 20x objective (0.45NA) were used to acquire images. Deconvolution of images acquired at the DeltaVision Spectris was performed using SoftWorx (Applied Precision).

### EdU assay

EdU (5-ethynyl-2’-deoxyuridine) was incorporated to cells as indicated in each experiment. A click-chemistry reaction was performed after the primary antibody washing step (See immunostaining procedure) using a mixture solution containing 100mM tris (pH 8.0), 1mM CuSO4, 100mM ascorbic acid and fluorescent dye (A10266 and A10277, Life technology), and incubated for 1 hour at room temperature.

### Correlative live-cell fluorescence and quantitative phase image (QPI) microscopy

Cells were seeded in a 6-well plate (Corning) at 4 x 10^4^ cells per well (approx. 30% confluency). Correlative fluorescence and QPI imaging was performed using the Livecyte microscope (Phase Focus). QPI and fluorescence images were acquired using a 10x objective (0.25 NA) every 10 and 30 mins, respectively for over 7 days, whilst being maintained at 37°C and 5% CO_2_. Cell culture media was replaced with fresh media every 3 days after the start of imaging. After each media exchange imaging was resumed of the same field of view. QPI and fluorescence images were exported as 8/16bit.tiff images using the Cell Analysis Toolbox software (Phase Focus) for subsequent analysis using ImageJ. For Figure 3b-d, cells were manually followed and scored. For Figure 3d-g, cells were manually tracked and nucleus and cytoplasm segmented based on intensity thresholds using a custom written macro. We selected to retain the cytoplasmic/nuclear thresholding for mitotic cells to provide a continuous read-out, giving rise to an arbitrary low Cdk activity measurement in mitotic cells. Cdk activity was calculated as the ratio of mean cytoplasmic and nuclear signal after rolling ball background subtraction. For QPI, pixel intensities in the exported.tiff images are proportional to dry mass, therefore integrated intensity after rolling ball background subtraction was used to represent total dry mass of the cell.

## Acknowledgements

This study has been supported by the Swedish Research Council and the Swedish Cancer Society (to AL). We thank Erik Müllers for initial observations and discussions.

## Supplementary data

Figure S1. Cell cycle progression after HU synchronisation release Cells harvested at 1,3,5,7,9 and 24 hours after HU (2mM) synchronisation release for cell cycle distribution analysis. The H bar indicates S-G2-M cell populations, the percentage of which is marked in each graph. The area in green indicates the total cell population, while the area in purple denotes the cyclin B1-eYFP positive population.

Movie S1. Combined Cdk activity sensor and QPI of cells shown in Figure 3. Images were cropped so that the tracked cells are in the middle of movies.

## References

1. Baus, F., Gire, V., Fisher, D., Piette, J. & Dulić, V. Permanent cell cycle exit in G2 phase after DNA damage in normal human fibroblasts. EMBO J. 22, 3992–4002 (2003).

2. Wiebusch, L. & Hagemeier, C. p53- and p21-dependent premature APC/C–Cdh1 activation in G2 is part of the long-term response to genotoxic stress. Oncogene 29, 3477–3489 (2010).

3. Gillis, L. D., Leidal, A. M., Hill, R. & Lee, P. W. K. p21Cip1/WAF1 mediates cyclin B1 degradation in response to DNA damage. Cell Cycle Georget. Tex 8, 253–256 (2009).

4. Lee, J., Kim, J. A., Barbier, V., Fotedar, A. & Fotedar, R. DNA Damage Triggers p21WAF1-dependent Emi1 Down-Regulation That Maintains G2 Arrest. Mol. Biol. Cell 20, 1891–1902 (2009).

5. de Boer, H. R., Guerrero Llobet, S. & van Vugt, M. A. T. M. Controlling the response to DNA damage by the APC/C-Cdh1. Cell. Mol. Life Sci. CMLS 73, 949–960 (2016).

6. Müllers, E., Silva Cascales, H., Jaiswal, H., Saurin, A. T. & Lindqvist, A. Nuclear translocation of Cyclin B1 marks the restriction point for terminal cell cycle exit in G2 phase. Cell Cycle Georget. Tex 13, 2733–2743 (2014).

7. Krenning, L., Feringa, F. M., Shaltiel, I. A., van den Berg, J. & Medema, R. H. Transient Activation of p53 in G2 Phase Is Sufficient to Induce Senescence. Mol. Cell 55, 59–72 (2014).

8. Johmura, Y. et al. Necessary and sufficient role for a mitosis skip in senescence induction. Mol. Cell 55, 73–84 (2014).

9. Feringa, F. M. et al. Persistent repair intermediates induce senescence. Nat. Commun. 9, 1–10 (2018).

10. Müllers, E., Silva Cascales, H., Burdova, K., Macurek, L. & Lindqvist, A. Residual Cdk1/2 activity after DNA damage promotes senescence. Aging Cell 16, 575–584 (2017).

11. Kirkland, J. L. & Tchkonia, T. Cellular Senescence: A Translational Perspective. EBioMedicine 21, 21–28 (2017).

12. Herranz, N. & Gil, J. Mechanisms and functions of cellular senescence. J. Clin. Invest. 128, 1238–1246 (2018).

13. Kuilman, T., Michaloglou, C., Mooi, W. J. & Peeper, D. S. The essence of senescence. Genes Dev. 24, 2463–2479 (2010).

14. Bartkova, J. et al. Oncogene-induced senescence is part of the tumorigenesis barrier imposed by DNA damage checkpoints. Nature 444, 633–637 (2006).

15. Feo, F. Preneoplastic Lesions. in Encyclopedia of Cancer (ed. Schwab, M.) 2977–2984 (Springer, 2011). doi:10.1007/978-3-642-16483-5_4724.

16. Significance of multiple mutations in cancer | Carcinogenesis | Oxford Academic. https://academic.oup.com/carcin/article/21/3/379/2365659.

17. Neurohr et al. - 2019 - Excessive Cell Growth Causes Cytoplasm Dilution An.pdf.

18. Storchova, Z. & Pellman, D. From polyploidy to aneuploidy, genome instability and cancer. Nat. Rev. Mol. Cell Biol. 5, 45–54 (2004).

19. Bunz, F. et al. Requirement for p53 and p21 to sustain G2 arrest after DNA damage. Science 282, 1497–1501 (1998).

20. Davoli, T., Denchi, E. L. & Lange, T. de. Persistent Telomere Damage Induces Bypass of Mitosis and Tetraploidy. Cell 141, 81–93 (2010).

21. Vaziri, C. et al. A p53-Dependent Checkpoint Pathway Prevents Re-replication. Mol. Cell 11, 997–1008 (2003).

22. Agarwal, M. L., Agarwal, A., Taylor, W. R. & Stark, G. R. p53 controls both the G2/M and the G1 cell cycle checkpoints and mediates reversible growth arrest in human fibroblasts. Proc. Natl. Acad. Sci. U. S. A. 92, 8493–8497 (1995).

23. p53: Guardian of ploidy - Aylon - 2011 - Molecular Oncology - Wiley Online Library. https://febs.onlinelibrary.wiley.com/doi/full/10.1016/j.molonc.2011.07.007.

24. Spencer, S. L. et al. The Proliferation-Quiescence Decision Is Controlled by a Bifurcation in CDK2 Activity at Mitotic Exit. Cell 155, 369–383 (2013).

25. Schwarz, C. et al. A Precise Cdk Activity Threshold Determines Passage through the Restriction Point. Mol. Cell 69, 253–264.e5 (2018).

26. Marrison, J., Räty, L., Marriott, P. & O’Toole, P. Ptychography – a label free, high-contrast imaging technique for live cells using quantitative phase information. Sci. Rep. 3, 2369 (2013).

27. Suman, R. et al. Label-free imaging to study phenotypic behavioural traits of cells in complex co-cultures. Sci. Rep. 6, 22032 (2016).

28. Fu, J., Hagan, I. M. & Glover, D. M. The centrosome and its duplication cycle. Cold Spring Harb. Perspect. Biol. 7, a015800 (2015).

29. Davoli, T. & de Lange, T. The Causes and Consequences of Polyploidy in Normal Development and Cancer. Annu. Rev. Cell Dev. Biol. 27, 585–610 (2011).

30. Shu, Z., Row, S. & Deng, W.-M. Endoreplication: The Good, the Bad, and the Ugly. Trends Cell Biol. 28, 465–474 (2018).

31. Edgar, B. A., Zielke, N. & Gutierrez, C. Endocycles: a recurrent evolutionary innovation for post-mitotic cell growth. Nat. Rev. Mol. Cell Biol. 15, 197–210 (2014).

32. Hsu, C.-H., Altschuler, S. J. & Wu, L. F. Patterns of Early p21 Dynamics Determine Proliferation-Senescence Cell Fate after Chemotherapy. Cell 178, 361–373.e12 (2019).

33. Yang, H. W., Chung, M., Kudo, T. & Meyer, T. Competing memories of mitogen and p53 signalling control cell-cycle entry. Nature 549, 404–408 (2017).

34. Barr, A. R. et al. DNA damage during S-phase mediates the proliferation-quiescence decision in the subsequent G1 via p21 expression. Nat. Commun. 8, 14728 (2017).

35. Arora, M., Moser, J., Phadke, H., Basha, A. A. & Spencer, S. L. Endogenous Replication Stress in Mother Cells Leads to Quiescence of Daughter Cells. Cell Rep. 19, 1351–1364 (2017).

36. Min, M., Rong, Y., Tian, C. & Spencer, S. Temporal integration of mitogen history in mother cells controls proliferation of daughter cells. Science (2020) doi:10.1126/science.aay8241.

37. Sherr, C. J. & Roberts, J. M. Inhibitors of mammalian G1 cyclin-dependent kinases. Genes Dev. 9, 1149–1163 (1995).

38. Potapova, T. A., Seidel, C. W., Box, A. C., Rancati, G. & Li, R. Transcriptome analysis of tetraploid cells identifies cyclin D2 as a facilitator of adaptation to genome doubling in the presence of p53. Mol. Biol. Cell 27, 3065–3084 (2016).

39. Reyes, J. et al. Fluctuations in p53 Signaling Allow Escape from Cell-Cycle Arrest. Mol. Cell 71, 581–591.e5 (2018).

40. Melixetian, M. et al. Loss of Geminin induces re-replication in the presence of functional p53. J. Cell Biol. 165, 473–482 (2004).

41. Sørensen, C. S. & Syljuåsen, R. G. Safeguarding genome integrity: the checkpoint kinases ATR, CHK1 and WEE1 restrain CDK activity during normal DNA replication. Nucleic Acids Res. 40, 477–486 (2012).

42. Lemmens, B. & Lindqvist, A. DNA replication and mitotic entry: A brake model for cell cycle progression. J. Cell Biol. 218, 3892–3902 (2019).

43. Matson, J. P. et al. Intrinsic checkpoint deficiency during cell cycle re-entry from quiescence. J. Cell Biol. 218, 2169–2184 (2019).

44. Zheng, L. et al. Fen1 mutations that specifically disrupt its interaction with PCNA cause aneuploidy-associated cancer. Cell Res. 21, 1052–1067 (2011).

45. Zheng, L. et al. Polyploid cells rewire DNA damage response networks to overcome replication stress-induced barriers for tumour progression. Nat. Commun. 3, 1–12 (2012).

46. Ganem, N. J., Godinho, S. A. & Pellman, D. A mechanism linking extra centrosomes to chromosomal instability. Nature 460, 278–282 (2009).

47. Sakaue-Sawano, A., Kobayashi, T., Ohtawa, K. & Miyawaki, A. Drug-induced cell cycle modulation leading to cell-cycle arrest, nuclear mis-segregation, or endoreplication. BMC Cell Biol. 12, 2 (2011).

48. Litwiniec, A., Gackowska, L., Helmin-Basa, A., Zuryń, A. & Grzanka, A. Low-dose etoposide-treatment induces endoreplication and cell death accompanied by cytoskeletal alterations in A549 cells: Does the response involve senescence? The possible role of vimentin. Cancer Cell Int. 13, 9 (2013).

49. Tan, Z., Chu, D. Z. V., Chan, Y. J. A., Lu, Y. E. & Rancati, G. Mammalian Cells Undergo Endoreduplication in Response to Lactic Acidosis. Sci. Rep. 8, 1–10 (2018).

50. Puig, P.-E. et al. Tumor cells can escape DNA-damaging cisplatin through DNA endoreduplication and reversible polyploidy. Cell Biol. Int. 32, 1031–1043 (2008).

51. Zhang, S. et al. Generation of cancer stem-like cells through the formation of polyploid giant cancer cells. Oncogene 33, 116–128 (2014).

52. Niu, N., Mercado-Uribe, I. & Liu, J. Dedifferentiation into blastomere-like cancer stem cells via formation of polyploid giant cancer cells. Oncogene 36, 4887–4900 (2017).

53. Akopyan, K. et al. Assessing Kinetics from Fixed Cells Reveals Activation of the Mitotic Entry Network at the S/G2 Transition. Mol. Cell 53, 843–853 (2014).

54. Cascales, H. S. et al. Cyclin A2 localises in the cytoplasm at the S/G2 transition to activate Plk1. bioRxiv 191437 (2017) doi:10.1101/191437.

