## Supplementary figures and images for "p53-dependent polyploidisation after DNA damage in G2 phase"

### Supplemental figure 1

#### S1 Cell cycle progression after HU synchronisation release

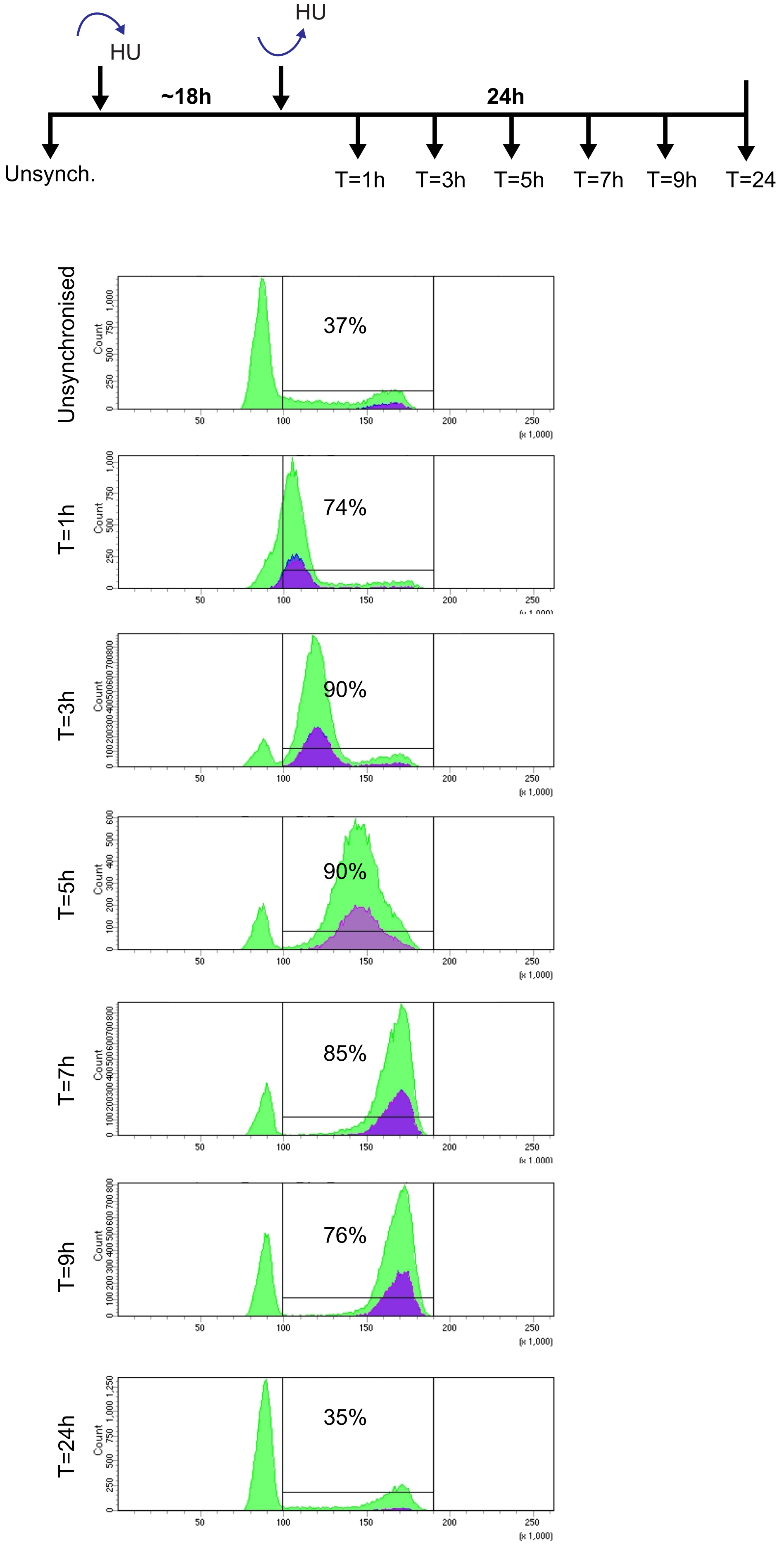
